# Epitranscriptomic *N*^6^-methyladenosine profile of SARS-CoV-2-infected human lung epithelial cells

**DOI:** 10.1101/2022.08.01.502311

**Authors:** Stacia Phillips, Shaubhagya Khadka, Dana Bohan, Constanza E. Espada, Wendy Maury, Li Wu

**Author notes:** Corresponding author. Li Wu.

## Abstract

*N*^6^-methyladenosine (m^6^A) is a dynamic post-transcriptional RNA modification that plays an important role in determining transcript fate. Severe acute respiratory syndrome-related coronavirus 2 (SARS-CoV-2) has caused the global pandemic of coronavirus disease 2019 (COVID-19) and the virus has been extensively studied. However, how m^6^A modification of host cell RNAs change during SARS-CoV-2 infection has not been reported. Here we define the epitranscriptomic m^6^A profile of SARS-CoV-2-infected human lung epithelial cells compared to uninfected controls. Biological pathway analyses revealed that differentially methylated transcripts were significantly associated with cancer-related pathways, protein processing in the endoplasmic reticulum, cell death and proliferation. Upstream regulators predicted to be associated with the proteins encoded by differentially methylated mRNAs include proteins involved in the type I interferon response, inflammation, and cytokine signaling. These data suggest that m^6^A modification of cellular RNA is an important mechanism of regulating host gene expression during SARS-CoV-2 infection of lung epithelial cells.

## Introduction

*N*^6^-methyladenosine (m^6^A) is the most prevalent post-transcriptional modification of eukaryotic mRNA and plays an important role in the fate the modified mRNA molecule. The m^6^A is deposited on adenosine by a methyltransferase, or “writer”, complex consisting of the catalytic heterodimer methyltransferase-like-3 and methyltransferase-like14 (METTL3/METTL14) in complex with the adapter protein Wilms tumor 1-associated protein (WTAP) (Liu et al., 2014). m^6^A is also prevalent on small non-coding RNA (sncRNA) and long non-coding RNA (lncRNA) and this modification is catalyzed by the writer METTL16 (Warda et al., 2017). Two demethylases, or “erasers”, fat mass and obesity-associated protein (FTO) and α-ketoglutarate-dependent dioxygenase AlkB homolog 5 (ALKBH5) can remove the m^6^A modification, suggesting that m^6^A modification is not only dynamic but reversible (Jia et al., 2011; Zheng et al., 2013). The outcome of m^6^A modification is dictated by m^6^A-specific RNA binding proteins or “readers”, the most well characterized of which are members of the YT521-B homology (YTH) family (Patil et al., 2018; Zaccara and Jaffrey, 2020). Binding of readers to the modified mRNA can lead to changes in stability, translation, localization, and splicing (Lesbirel and Wilson, 2019; Roundtree et al., 2017; Wang et al., 2014; Wang et al., 2015; Zhao et al., 2014; Zheng et al., 2017). Therefore, m^6^A modification acts as an important mechanism of post-transcriptional regulation of gene expression.

Many virus genomes and viral RNAs are m^6^A-modified, and these modifications play important functional roles in various stages of virus replication and evasion of innate immune sensing (Imam et al., 2020). SARS-CoV-2 RNAs are m^6^A-modified and while some cell type-dependent discrepancies exist, most studies have reported that m^6^A is required for efficient virus replication (Burgess et al., 2021; Campos et al., 2021; Li et al., 2021a; Liu et al., 2021; Zhang et al., 2021). In addition to functional m^6^A modification of viral RNAs, changes in the cellular m^6^A methylome have also been shown to occur in association with viral infections (Gokhale et al., 2020; McFadden and Horner, 2021; Williams et al., 2019). Of particular interest, cellular transcripts involved in establishing an antiviral immune response are post-transcriptionally regulated by m^6^A modification (McFadden et al., 2021; Rubio et al., 2018; Winkler et al., 2019). It is likely that SARS-CoV-2 infection leads to changes in the m^6^A modification state of host cell transcripts, either induced directly by the virus or through the cellular response to infection. Indeed, m^6^A sequencing (m^6^A-seq or meRIP-seq) has revealed the loss or gain of m^6^A modifications in host cell RNA from infected cells (Liu et al., 2021). However, how the host m^6^A methylome changes in human lung cells in response to SARS-CoV-2 infection remains unknown.

Here we report the results of epitranscriptomic m^6^A microarray analysis of human lung cells infected with SARS-CoV-2 compared to uninfected control cells. We identified changes in the abundance of methylated cellular RNAs for both protein-coding and non-coding transcripts. One micro-RNA (miR) precursor, miR-4486, was found to be 175 times more abundant in the methylated fraction of infected-cell RNA compared to uninfected controls. Interestingly, biological pathway analysis revealed that many differentially methylated mRNA transcripts code for proteins that are regulated upstream by proteins involved in inflammation, cytokine signaling, and innate immunity. These findings will serve as the basis for future functional validation studies to determine how changes in the methylation status of host cell transcripts may affect SARS-CoV-2 replication and viral pathogenesis.

## Results

### A549-hACE2 cells support productive SARS-CoV-2 infection

We sought to determine the epitranscriptomic m^6^A profile of SARS-CoV-2-infected cell RNA using a human lung epithelial cell line, as lung epithelial cells represent a biologically relevant target of SARS-CoV-2. Robust and reliable identification of changes to the methylation level of individual host cell transcripts during infection is best achieved using conditions under which most of the cells have become infected, reducing background signal contributed by uninfected cells. Therefore, we first directly compared three different lung cell lines (A549-hACE2 cells expressing human angiotensin-converting enzyme 2 [hACE2], Calu-3, and H1650) for their ability to support SARS-CoV-2 replication under identical conditions. In our infection assays, we chose to infect cells with SARS-CoV-2 (strain USA-WA-1/2020) at a multiplicity of infection (MOI) of 1 for 24 hours to allow for a full viral life cycle and spreading infection to occur (Li et al., 2021b). After 24 hours, A549-hACE2 cells were fixed, and infected cells were visualized by immunofluorescent staining using a SARS-CoV-2 nucleocapsid-specific antibody (Fig. 1A). Infection of A549-hACE2 cells resulted in a greater proportion of N-positive cells (∼70%) compared to Calu-3 and H1650 cell lines (data not shown). Real-Time quantitative PCR (RT-qPCR) analysis using *spike* gene-specific qPCR primers demonstrated robust viral RNA replication in A549-hACE2 cells with ∼5 × 10^4^ copies of spike RNA present per infected cell (Fig. 1B). These RNA molecules represent both full-length positive sense RNA and subgenomic RNA used for translation to viral protein. Based on these results, we chose to use A549-hACE2 cells to determine the epitranscriptomic m^6^A profile of SARS-CoV-2-infected cells.

**Fig. 1.**
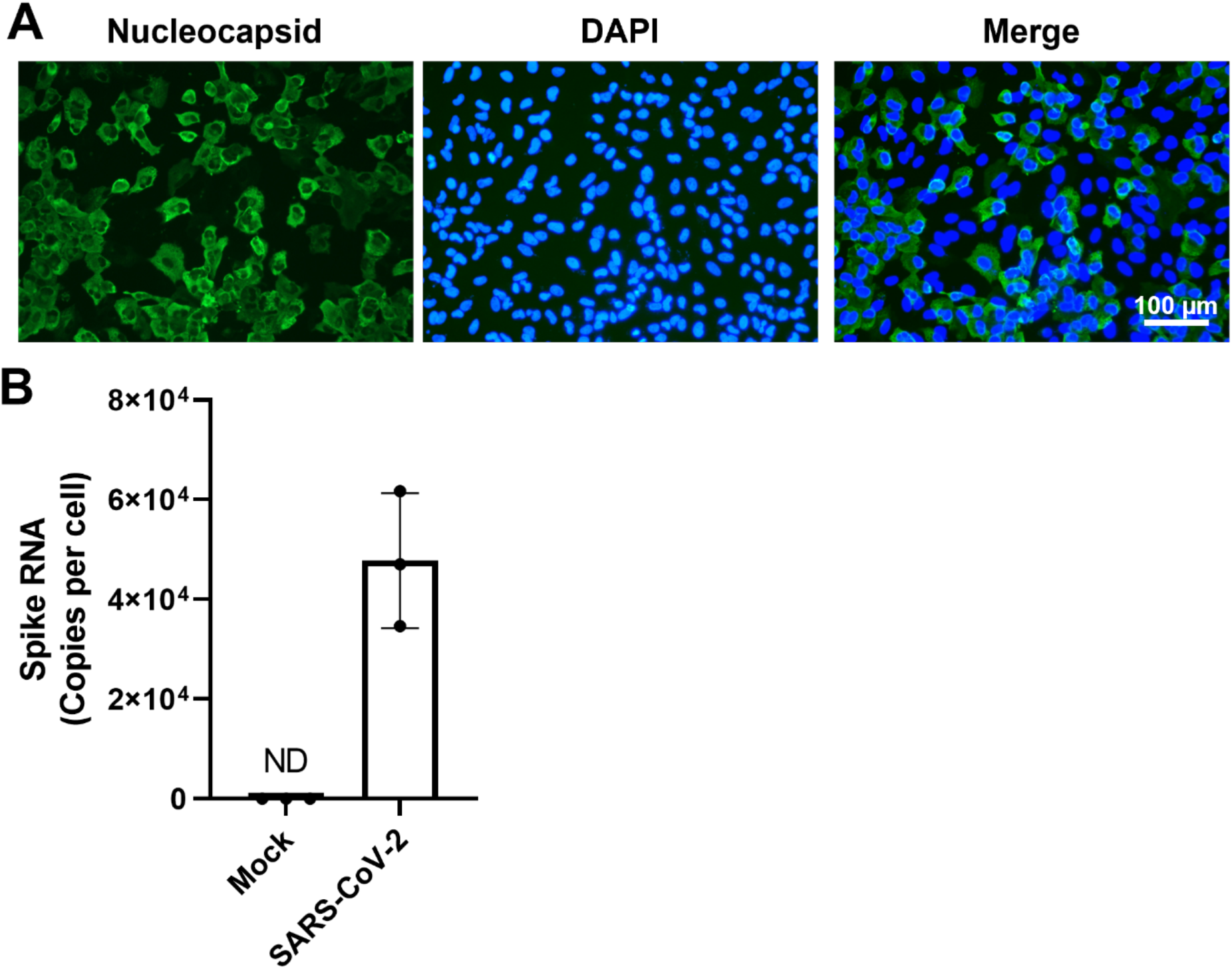
SARS-CoV-2 infection of A549-hACE2 cells. A549-hACE2 cells were infected with SARS-CoV-2 (strain USA-WA1/2020) at an MOI of 1 for 24 hours. (**A**) Immunofluorescent staining was performed to visualize infected cells by the presence of SARS-CoV-2 nucleocapsid (green). Nuclei of cells are stained with DAPI (blue). (**B**) SARS-CoV-2 spike RNA in infected cells (N=3, biological triplicate) was quantified by RT-qPCR. ND: not detected.

### SARS-CoV-2 infection of A549-hACE2 cells leads to differential m^6^A modification of cellular RNA

To analyze m^6^A modifications of host cell transcripts, A549-hACE2 cells were infected with SARS-CoV-2 at an MOI of 1 for 24 hours in biological triplicate. Total RNA from SARS-CoV-2-infected cells and mock-infected negative control cells was used for m^6^A immunoprecipitation (IP) followed by microarray analysis using the Arraystar Epitranscriptomic m^6^A Array (see schematic method summary in Fig. 2A). The RNA present in the IP fraction represents m^6^A-modified RNA, whereas the remaining unbound fraction is assumed to be unmethylated. Each fraction was fluorescently labeled and mixed prior to array hybridization. The microarray contains over 60,000 unique probes (60 nt each) that represent 44,122 mRNA, 12,496 lncRNA, 1,366 pre-miRNA, 1,642 pri-miRNA, 19 small nuclear RNA (snRNA), and 786 small nucleolar RNA (snoRNA) transcripts. Unique mRNA splice isoforms are distinguished by probes that are exon specific or span a splice junction. Signal for each transcript in the IP and unbound fractions was normalized to the intensity of non-human spike-in RNA.

**Fig. 2.**
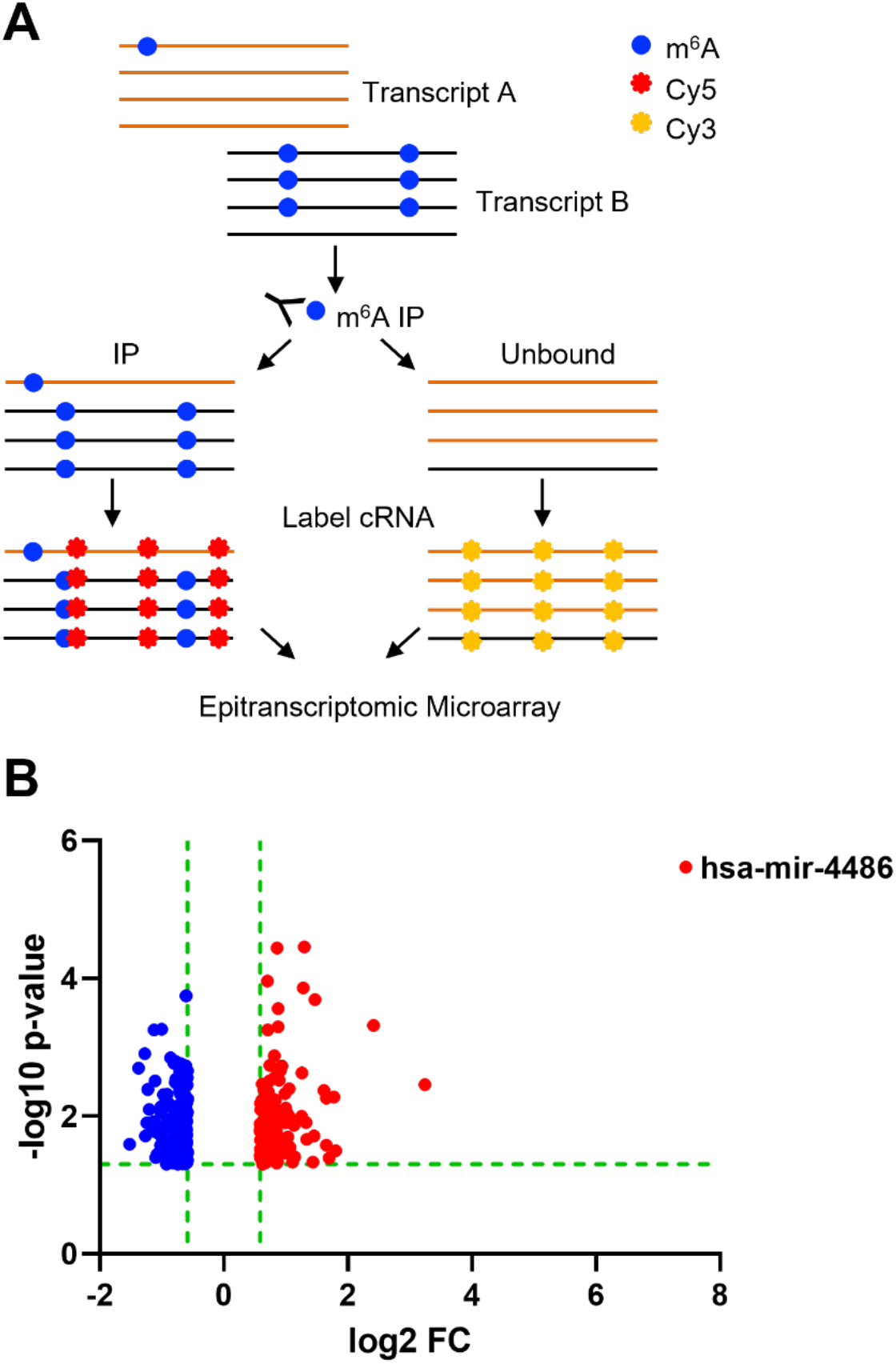
Epitranscriptomic m^6^A microarray of SARS-CoV-2-infected A549-hACE2 cells. (**A**) Schematic overview of the method. Total cellular RNA from each sample (SARS-CoV-2-infected and mock-infected controls, biological triplicate, N=3 each group) was used for immunoprecipitation using an m^6^A-specific antibody. Methylated and unmethylated RNA fractions were fluorescently labeled (Cy3 or Cy5) prior to array hybridization (refer to Materials and Methods for details). (**B**) Volcano plot of transcripts containing higher (red) and lower (blue) levels of m^6^A modification in infected cells compared to mock-infected control cells. The miRNA precursor (hsa-mir-4486) with the most significant m^6^A change is labeled.

Transcript types that were found to be significantly differentially methylated ≥ 1.5-fold with a p-value ≤ 0.05 in SARS-CoV-2-infected cells compared to mock-infected controls are summarized in Table 1. A total of 186 unique transcripts were hypomethylated and 119 transcripts were hypermethylated in response to infection. A volcano plot shows the statistical significance (-log10 p-value) versus the fold change (log2 FC) in the abundance of methylated transcript in SARS-CoV-2-infected cells relative to mock (Fig. 2B). A selection of transcripts with the most significant changes in m^6^A abundance (p-value ≤ 0.005) are shown in Table 2. The full list of transcripts with significantly different m^6^A abundance in infected cells can be found in Table S1 (p-value ≤ 0.05, fold change ≥ 1.5).

**Table 1.** Summary of differentially modified transcripts by type in SARS-CoV-2-infected A549-hACE2 cells vs. mock-infected control cells.

| Transcript Type | Hypo m<sup>6</sup>A | Hyper m<sup>6</sup>A | Total | Numbers of Probes |
| --- | --- | --- | --- | --- |
| mRNA | 160 | 89 | 249 | 44,122 |
| lncRNA | 24 | 22 | 46 | 12,496 |
| sncRNA | 2 | 8 | 10 | 3,813 |

Fold change $\geq 1.5$ and $p \leq 0.05$

**Table 2.** Selected differentially m^6^A-modified transcripts in SARS-CoV-2-infected A549-hACE2 cells vs. mock-infected control cells by m^6^A quantity.

| Transcript Type | Transcript ID | Gene Symbol | Log2 Fold Change | p-value |
| --- | --- | --- | --- | --- |
| mRNA | ENST00000250457 | EGLN3 | 0.8595436 | 3.609E-05 |
| mRNA | HBMT00001402195 | CATG00000101027.1 | 1.4687341 | 0.0002032 |
| mRNA | ENST00000540159 | BNIP3 | 0.8732292 | 0.0002728 |
| mRNA | ENST00000325885 | ASB4 | 2.4125726 | 0.0004796 |
| mRNA | ENST00000618819 | PTPN9 | 0.8113196 | 0.0013265 |
| mRNA | NM_001330464 | ATL2 | 0.9303607 | 0.0018649 |
| mRNA | ENST00000357077 | ANK2 | 0.8673231 | 0.0021318 |
| mRNA | ENST00000422053 | TRIB3 | 0.9064073 | 0.0021784 |
| mRNA | ENST00000528331 | SYBU | 1.255638 | 0.0023711 |
| mRNA | ENST00000311450 | PLAC8L1 | 0.8936838 | 0.0029927 |
| mRNA | ENST00000382181 | RBCK1 | 0.7350803 | 0.0030862 |
| mRNA | ENST00000248071 | KLF2 | 0.6192452 | 0.0034099 |
| mRNA | ENST00000374448 | MUSK | 3.2415767 | 0.0034892 |
| mRNA | ENST00000376468 | NPPB | 1.0536428 | 0.0040088 |
| mRNA | ENST00000371445 | DMRTB1 | 0.6458079 | 0.0041402 |
| mRNA | ENST00000644112 | TIMM8A | -0.6091437 | 0.0001784 |
| mRNA | ENST00000360948 | NTRK3 | -1.0030983 | 0.0005479 |
| mRNA | ENST00000629765 | NTRK3 | -1.1272045 | 0.0005579 |
| mRNA | ENST00000394480 | NTRK3 | -1.2754039 | 0.0012382 |
| mRNA | ENST00000417456 | RP11-244H3.5 | -0.8596983 | 0.0014264 |
| mRNA | ENST00000619416 | KIAA0586 | -0.7813802 | 0.0015997 |
| mRNA | ENST00000539097 | HIF1A | -0.8053586 | 0.0017167 |
| mRNA | ENST00000451528 | ST8SIA4 | -0.6902188 | 0.0017212 |
| mRNA | NM_001320134 | NTRK3 | -0.61504 | 0.0018528 |
| mRNA | ENST00000378133 | PCDHA1 | -0.5856729 | 0.0022007 |
| mRNA | ENST00000333896 | SPTBN1 | -0.7070035 | 0.002238 |
| mRNA | ENST00000368581 | RSPH4A | -0.6288855 | 0.0026367 |
| mRNA | ENST00000394332 | RPL23 | -0.6408456 | 0.0026379 |
| mRNA | ENST00000357496 | QRICH1 | -0.593256 | 0.0027733 |
| mRNA | ENST00000537259 | SLC24A1 | -0.7748985 | 0.0028879 |
| mRNA | MICT00000118072 | CATG00000023300.1 | -0.6383486 | 0.0029291 |
| mRNA | ENST00000644112 | TIMM8A | -0.6091437 | 0.0001784 |
| mRNA | ENST00000360948 | NTRK3 | -1.0030983 | 0.0005479 |
| lncRNA | NR_148507 | AC010982.1 | 1.2975579 | 3.476E-05 |
| lncRNA | ENST00000477817 | PKP1 | 0.7016014 | 0.0001088 |
| lncRNA | NR_152836 | PSORS1C3 | 1.2784829 | 0.0001375 |
| lncRNA | ENST00000579368 | RP11-674N23.1 | 0.8775907 | 0.0005063 |
| lncRNA | ENST00000592680 | AC007773.2 | 0.7066193 | 0.0005585 |
| lncRNA | ENST00000436551 | AC104654.2 | -1.3765495 | 0.0020037 |
| lncRNA | NR_136184 | CENPO | -0.7059265 | 0.0026148 |
| lncRNA | ENST00000503936 | TTC33 | -0.6501871 | 0.0027399 |
| lncRNA | ENST00000422847 | RP11-40F6.2 | -0.6267391 | 0.0050007 |
| pre-miRNA | MI0001446 | hsa-mir-4486 | 7.4490146 | 2.413E-06 |

p ≤ 0.005

The plot of log2 FC shows that changes in hypomethylated transcripts ranged from a log2 FC of -0.59 to -1.52 (1.5-fold to 2.8-fold change). Similarly, the log2 FC for most transcripts found to be hypermethylated in infected cells ranged from 0.59 to 1.47 (1.5-fold to 2.8-fold change). Eight of the hypermethylated transcripts showed a greater than 3-fold change. Remarkably, the primary miR transcript for miR-4486 was found in the methylated RNA fraction from infected cells at a 175-fold higher level than in uninfected controls. The associated p-value of this change was 2.41 × 10^−6^, demonstrating high reproducibility among three independent infections and uninfected controls (Fig. 2B and Table S1).

A unique feature of the epitranscriptomic microarray is the ability to determine the percentage of transcript molecules that are m^6^A-modified, based on the relative intensity of signals in the IP and unbound fractions. This stoichiometric information is not provided by other m^6^A detection techniques such as m^6^A-seq (Dominissini et al., 2012; McIntyre et al., 2020). Selected transcripts with a significantly different percentage of m^6^A-modified RNA in SARS-CoV-2-infected samples relative to mock controls are shown in Table 3 (p-value ≤ 0.005). The full list of transcripts with a significantly different percentage of m^6^A-modified RNA in infected cells can be found in Table S1 (p-value ≤ 0.05). These results show that many cellular transcripts undergo changes in m^6^A abundance and percentage of transcript modified in response to SARS-CoV-2 infection of A549-hACE2 cells.

**Table 3.** Selected differentially m^6^A-modified transcripts in SARS-CoV-2-infected A549-hACE2 cells vs. mock-infected control cells by % modified.

| Transcript Type | Transcript ID | Gene Symbol | % Mock | % SARS-CoV-2 |
| --- | --- | --- | --- | --- |
| mRNA | ENST00000374448 | MUSK | 30% | 69% |
| mRNA | ENST00000371968 | LHFPL1 | 33% | 43% |
| mRNA | ENST00000264657 | STAT3 | 27% | 34% |
| mRNA | ENST00000528331 | SYBU | 40% | 53% |
| mRNA | ENST00000360375 | LRRCC1 | 40% | 23% |
| mRNA | ENST00000361987 | CNTF | 91% | 81% |
| mRNA | ENST00000216513 | SIX4 | 93% | 82% |
| mRNA | ENST00000367512 | EDEM3 | 83% | 69% |
| mRNA | ENST00000522232 | CTB-83D3.1 | 72% | 34% |
| mRNA | ENST00000361028 | ZSCAN12 | 80% | 54% |
| lncRNA | NR_145484 | ARHGAP32 | 91% | 74% |
| lncRNA | NR_015352 | CECR7 | 65% | 32% |
| pre-miRNA | MI0001446 | hsa-mir-4486 | 44% | 73% |

p ≤ 0.005

### Biological pathway analysis of differentially methylated protein-coding transcripts

Protein coding transcripts were analyzed by iPathwayGuide (Advaita) to identify biological pathways that are significantly associated with cellular mRNAs that are differentially methylated in response to SARS-CoV-2 infection of A549-hACE2 cells. The p-values for significance of the association are derived from a combination of two independent analyses, classical over-representation analysis (pORA) and a measure of accumulated perturbation of a given pathway (pAcc) (Ahsan and Draghici, 2017; Donato et al., 2013; Draghici et al., 2007; Tarca et al., 2009). The full list of pathways associated with at least one differentially methylated mRNA is listed in Table S2, with pAcc, pORA, and the combined p-value indicated. Pathways with no pAcc represent metabolic networks, as opposed to signaling pathways which have both pAcc and pORA.

Fig. 3 shows the list of pathways with the highest combined significance (p ≤ 0.005). We found that protein coding transcripts that are differentially methylated in response to SARS-CoV-2 infection are associated with several cancer-related pathways (microRNAs, pathways, proteoglycans, and programmed death ligand 1 [PD-L1] expression and PD-1 checkpoint pathways in cancer), infectious disease (Legionellosis, Kaposi sarcoma-associated herpesvirus infection, and Hepatitis B), cell metabolism, proliferation, and survival/death (protein processing in the endoplasmic reticulum, metabolic pathways, necroptosis, forkhead box O [FoxO] signaling, mitophagy, epidermal growth factor receptor [EGFR] tyrosine kinase inhibitor resistance, signaling pathways regulating pluripotency of stem cells, phosphoinositide-3-kinase-Akt [PI3K-Akt] signaling), and the immune response (JAK-STAT signaling). Our data also indicated the number of differentially methylated transcripts that are associated with each pathway (count) and the -log10 of the associated combined p-value. Together, these results suggest complex and dynamic biological pathways are involved in cellular responses to SARS-CoV-2 infection.

**Fig. 3.**
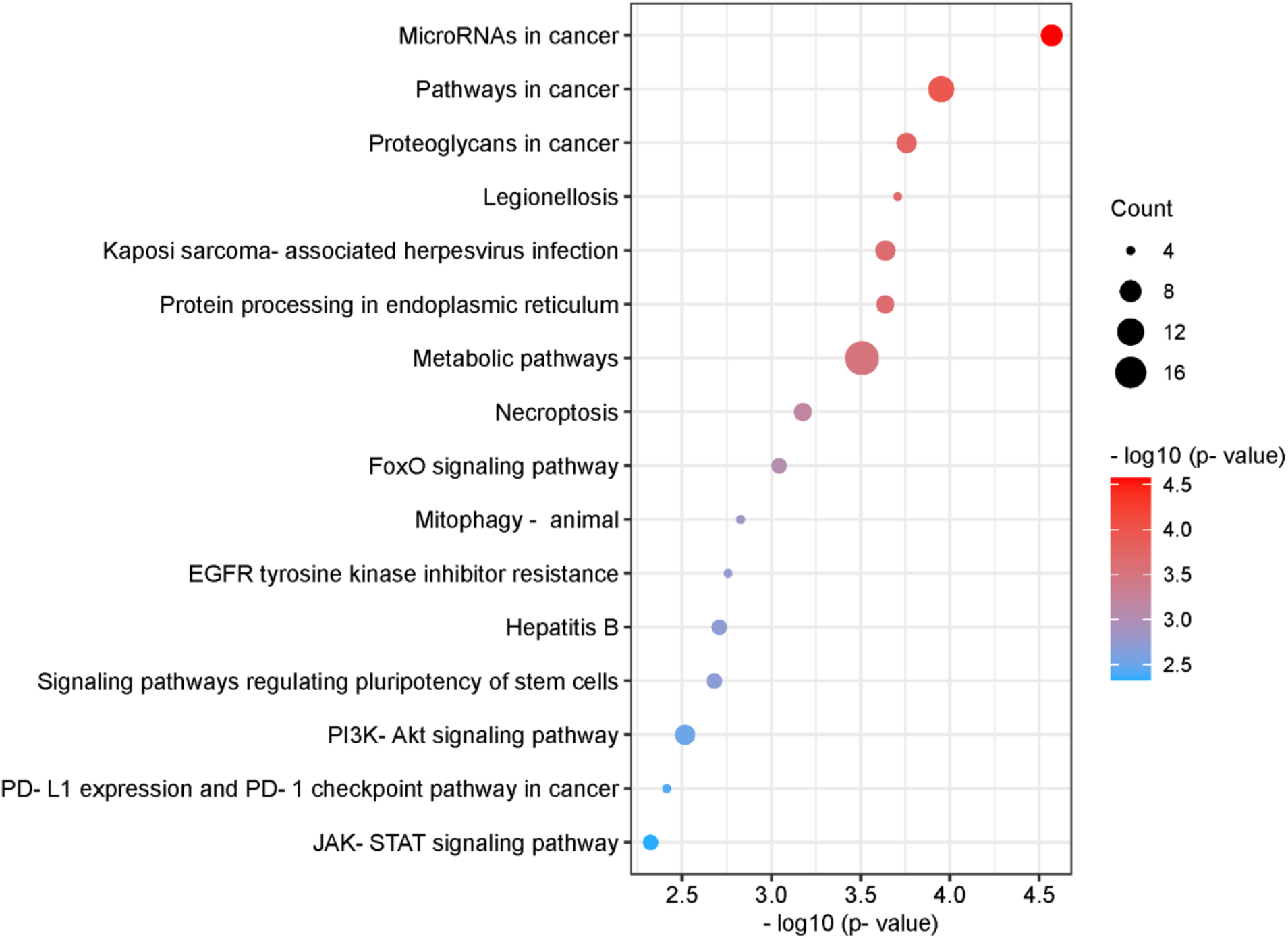
Pathway analysis of differentially methylated mRNAs. List of pathways associated with differentially methylated mRNAs with the highest combined significance (pORA and pAcc ≤ 0.005). Significance is indicated on the x-axis and by sphere color. The size of the sphere corresponds to the number of differentially methylated mRNAs associated with each pathway (count). Bubble plot was created using https://www.bioinformatics.com.cn/en, a free online platform for data analysis and visualization.

### Upstream regulators of differentially methylated mRNAs and predicated networks

To further analyze the significant regulators of differentially methylated transcripts during SARS-CoV-2 infection, iPathwayGuide was also used to identify putative upstream regulators of proteins encoded for by transcripts found to be differentially methylated in SARS-CoV-2-infected cells compared to mock-infected control cells. Interestingly, the top 20 upstream regulators predicted with the highest significance (p ≤ 0.01) were enriched for proteins involved in inflammation, cytokine signaling, and innate immunity (Fig. 4). The most significantly associated predicted upstream regulator is EGFR, which has been implicated as a potential therapeutic target for COVID-19 treatment (Klann et al., 2020; Londres et al., 2022; Vagapova et al., 2021; Venkataraman and Frieman, 2017). Other predicted upstream regulators with known function in inflammation and innate immunity include mitogen-activated protein kinase kinase 7 (MAP2K7), tumor necrosis factor receptor superfamily member 1A (TNFRSF1A), sprouty RTK signaling antagonist 4 (SPRY4), Janus kinase 3 (JAK3), Janus kinase 2 (JAK2), interferon alpha 1 (IFNA1), and tumor necrosis factor (TNF).

**Fig. 4.**
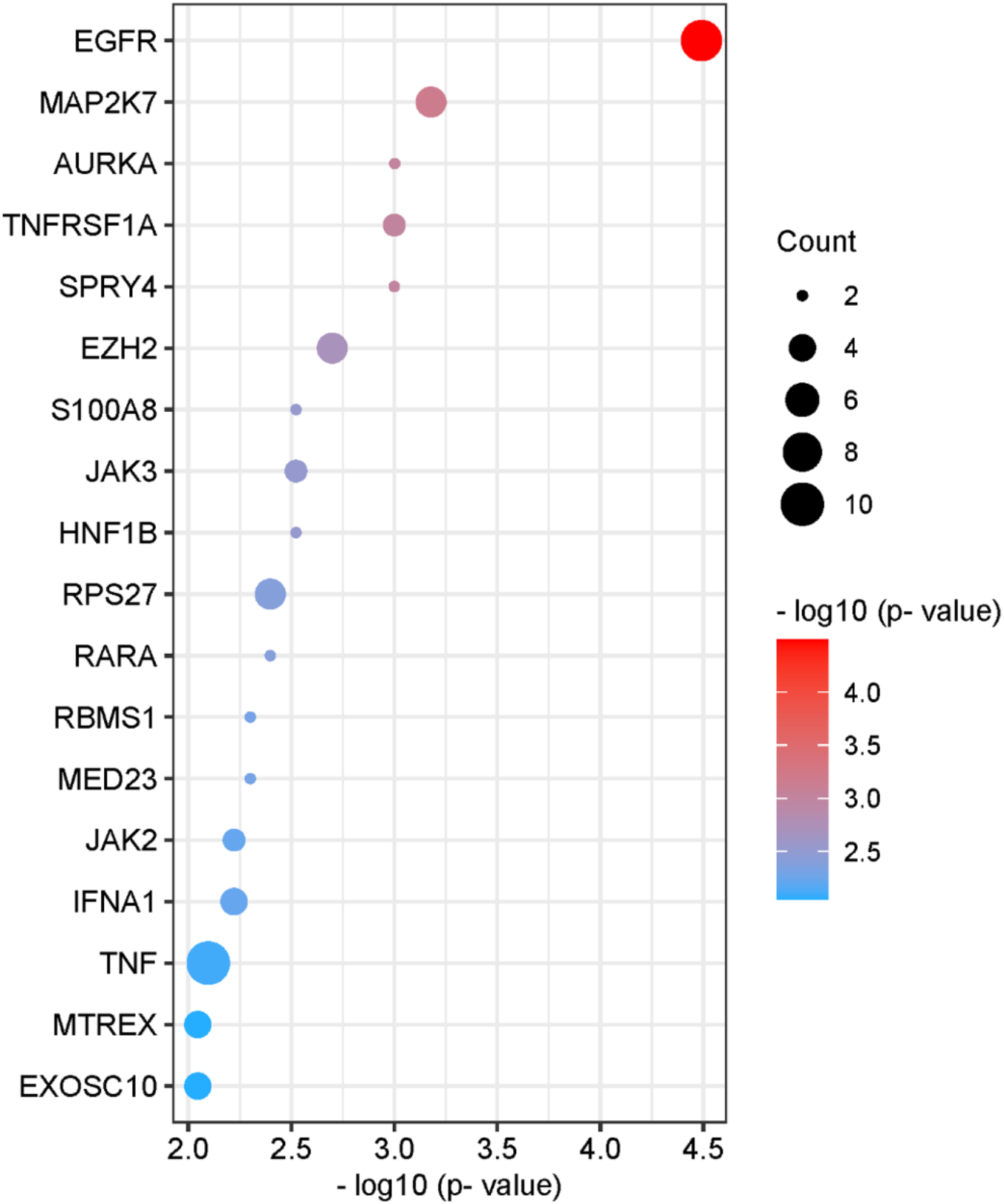
Upstream regulators of differentially methylated mRNAs. List of upstream regulators associated with differentially methylated mRNAs with the highest significance (p ≤ 0.01). Significance is indicated on the x-axis and by sphere color. The size of the sphere corresponds to the number of differentially methylated mRNAs associated with each pathway (count). Bubble plot was created using https://www.bioinformatics.com.cn/en, a free online platform for data analysis and visualization.

To better understand the interactions among significant regulators, predicted upstream regulators of differentially methylated transcripts were used to construct networks illustrating known regulatory interactions among individual nodes of a given pathway. Selected upstream regulators and their downstream targets that are differentially methylated in response to SARS-CoV-2 infection are indicated, with hypomethylated transcripts shown in blue and hypermethylated transcripts in pink (Fig. 5A-C). Gray nodes represent intermediate genes in the pathway that directly regulate or are regulated by differentially methylated pathway members, but do not themselves exhibit any change in methylation status upon SARS-CoV-2 infection. Pink arrows illustrate activation and gray bars represent inhibition for functional interactions that have been experimentally validated (Fig. 5A-C). These analyses allow us to develop novel hypotheses regarding how the host cell responds to SARS-CoV-2 infection. Together, these results suggest that regulation of gene expression at the level of post-transcriptional RNA modification is a mechanism by which the cell responds to SARS-CoV-2 infection and may have effects on viral pathogenesis and the immune response.

**Fig. 5.**
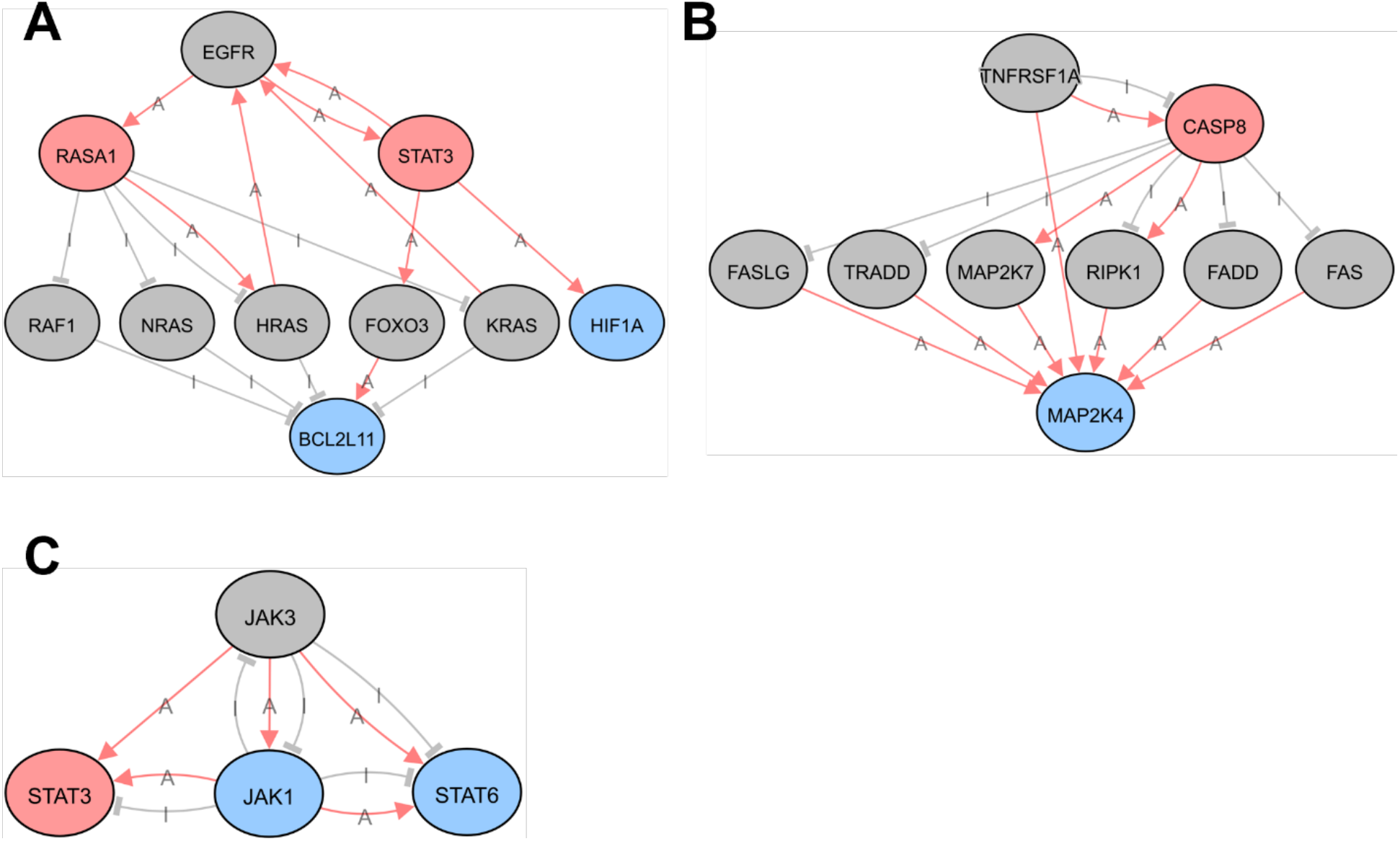
Predicted upstream regulator networks. Network map of selected upstream regulators of differentially methylated mRNA transcripts. Upstream regulators are **(A)** EGFR, **(B)** TNFRSF1A, and **(C)** JAK3. Colors represent the change in m^6^A modification of indicated transcripts in response to SARS-CoV-2 infection in human lung epithelial cells. Gray circles: no change; pink circles: hypermethylated; blue circles: hypomethylated. Lines indicate known functional interactions between pathway nodes. Pink arrows: activation (A); gray bars: inhibition (I). Images were obtained using iPathwayGuide from AdvaitaBio.

## Discussion

Post-transcriptional m^6^A modification of RNA is an important strategy for regulation of gene expression (Shi et al., 2019). We sought to identify changes in m^6^A modification of cellular RNA during SARS-CoV-2 infection of human lung epithelial cells using epitranscriptomic m^6^A microarray analysis. These cellular RNAs may be important for virus replication or for establishing an antiviral innate immune response. We identified mRNA and long and small non-coding RNA species that are differentially m^6^A-modified in response to SARS-CoV-2 infection. Differentially methylated mRNA transcripts were found to be associated with biological pathways and upstream regulators that are involved in the immune response to viral infection. These data may provide a basis for novel hypotheses regarding mechanisms of SARS-CoV-2 replication or the cellular response to infection of lung epithelial cells. Future functional studies of the identified cellular RNA are required to test these hypotheses.

Overall, we observed differential methylation of 305 unique transcripts, with 186 hypomethylated and 119 hypermethylated transcripts in infected cells compared to uninfected controls. These differentially methylated transcripts are a relatively small percentage of all transcripts represented on the epitranscriptomic microarray (Table 1). The number of transcripts and magnitude of differential methylation in the current study is likely an underestimation of the actual change in methylation status that occurs in infected cells, due to the contribution of RNA from the ∼30% of cells in infected cultures that remained uninfected when the RNA was harvested (Fig. 1A). Some effects may also be missed due to differences in IP efficiency of individual transcripts, due either to RNA length or location of the m^6^A modification(s). Although the consensus RRACH motif (R = A or G; H = A, C, or U) for m^6^A modification is relatively common, only ∼5% of these motifs are m^6^A modified, resulting in an average of 1-2 m^6^A modifications per mRNA transcript (Dominissini et al., 2012). Therefore, despite the limitations described above, it is possible that modest changes in m^6^A modification can potentiate changes to RNA function during SARS-CoV-2 infection.

One mRNA found to be significantly more m^6^A-modified in infected cells compared to mock-infected controls is the muscle-associated receptor tyrosine kinase (MUSK). The epitranscriptomic microarray indicated that this mRNA is present at ∼8-fold greater abundance in the m^6^A-IP fraction in infected cells and that the percentage of methylated transcript increased from 30% in uninfected cells to 69% in SARS-CoV-2-infected cells (Tables 2 and 3). MUSK is a receptor tyrosine kinase that is essential for the formation and maintenance of the neuromuscular junction and is expressed at very low levels in the lung under normal conditions (Uhlen et al., 2015). Auto-antibodies directed against MUSK inhibit acetylcholine receptor clustering at the neuromuscular junction and are associated with a rare form of myasthenia gravis (MG), a chronic autoimmune disorder in which antibodies destroy the communication between nerves and muscle, resulting in weakness of the skeletal muscles (Hoch et al., 2001). Interestingly, a recent case report identified the development of MUSK-associated MG potentially triggered by SARS-CoV-2 infection (Assini et al., 2021). It would be important to investigate how SARS-CoV-2 infection could induce the development of autoantibodies to MUSK or how this might involve post-transcriptional m^6^A modifications of the MUSK mRNA.

We also observed a modest but significant increase in the levels of m^6^A modification of signal transducer and activator of transcription-3 (STAT3) transcript (Table 3). This transcription factor plays a pivotal role in intracellular signaling and subsequent activation of gene expression in response to a variety of cytokines and chemokines, including IL-6 and type I interferons (Villarino et al., 2017). STAT3 may contribute to the pathogenesis of SARS-CoV-2 infection in a variety of ways considering its pleiotropic effects on inflammation and the immune response [reviewed in (Jafarzadeh et al., 2021)]. We also observed changes in the abundance of m^6^A-modified JAK1 and STAT6 transcripts, which are also part of the JAK/STAT signaling pathway and may functionally interact with each other as part of the host cell response to infection (Table S1 and Fig. 5C). Consistent with these results, pathway analysis revealed that the JAK-STAT pathway was significantly associated with differentially methylated mRNAs in infected cells (Fig. 3). Therefore, other members of the JAK-STAT pathway were identified as upstream regulators of the protein products of mRNAs found to be differentially methylated in infected cells (Fig. 4, Fig. 5A and 5C). This overrepresentation of the JAK-STAT pathway in our analyses may reflect the activation of the JAK-STAT pathway in both bystander and infected cells.

The cellular RNA found to be the most significantly differentially methylated in response to SARS-CoV-2 infection was the precursor of miR-4486 (Tables 2 and 3, and Fig. 2B). Due to the size of the microarray probes (60 nt), miR transcripts are represented by the unprocessed primary and precursor transcripts, which may serve as a proxy for the mature 22 nt functional miR. One functionally validated target of miR-4486 is JAK3, which was also identified in our analysis as an upstream regulator significantly associated with differentially methylated transcripts in SARS-CoV-2-infected cells (Figs. 4 and 5C) (Zhou et al., 2022). One possible hypothesis based on our network analysis is that degradation of JAK3 transcript by miR-4486 leads to lower STAT3 activation as a compensatory mechanism in the infected cell to counteract SARS-CoV-2-induced STAT3 hyperactivation (Fig. 5C) (Matsuyama et al., 2020). The cellular lncRNA SNHG20 acts as a competing endogenous RNA to sponge miR-4486 and prevent degradation of miR-4486 target transcripts (Liu et al., 2019; Liu et al., 2022). Interestingly, a meta-analysis of transcriptomic data sets from COVID-19 patient samples found that SNHG20 was among the top 10 most significantly upregulated lncRNAs (Chakraborty et al., 2021). It is possible that miR-4486 is upregulated early in infected cells, whereas SNHG20 is upregulated at later times to counteract inhibition of cellular gene expression by miR-4486. Finally, experimentally validated binding targets of miR-4486 include TNF receptor-associated factor 7 (TRAF7) (Karginov and Hannon, 2013) and interleukin 1 receptor-associated kinase 3 (IRAK3) (Karginov and Hannon, 2013), both of which are involved in innate immune signaling and can lead to inhibition of NF-κB activation (Kobayashi et al., 2002; Zotti et al., 2012). Further studies are needed to determine which of these miR-4486 targets are functionally relevant during SARS-CoV-2 infection of human lung epithelial cells.

EGFR was identified as the putative upstream regulator predicted to be associated with differentially methylated mRNA transcripts with the highest confidence (Fig. 4). A network map illustrates the potential functional interactions between EGFR and proteins downstream whose mRNA were found to be hypermethylated (RAS p21 protein activator 1 [RASA1] and STAT3) and hypomethylated (hypoxia inducible factor 1 [HIF1A] and BCL2 like 11 [BCL2L11]). Several studies have demonstrated that EGFR is highly expressed and that EGFR signaling activity contributes to lung fibrosis in COVID-19 patients, leading to the identification of EGFR as a potential therapeutic target for treating severe COVID-19 (Klann et al., 2020; Londres et al., 2022; Vagapova et al., 2021).

The effect of reversible and dynamic m^6^A modification on a given transcript is context dependent and may be inhibitory (destabilization or sequestration) or activating (enhanced translation or splicing) (Lesbirel and Wilson, 2019; Roundtree et al., 2017; Wang et al., 2014; Wang et al., 2015; Zhao et al., 2014; Zheng et al., 2017). Networks such as those shown in Fig. 5 are useful for predicting functional associations between differentially methylated transcripts and identifying other members of a given signaling pathway that may also be affected by the change in m^6^A modification. However, due to potential opposing functional effects of m^6^A modification on an mRNA, it may not be straightforward to predict the directionality of effect on a downstream pathway member caused by changes in methylation of an upstream mRNA target.

In summary, our analysis identifies many cellular RNAs that exhibit differential methylation in SARS-CoV-2-infected human lung epithelial cells. These results can lay the foundation for the broader research community for the formation of novel hypotheses regarding the role of post-transcriptional regulation of host gene expression during SARS-CoV-2 infection. Our future studies are focused on functional validation of selected transcripts and determining the biological significance of differential methylation of specific transcripts during SARS-CoV-2 infection of primary human airway epithelia cells.

## Material and methods

### Cells and SARS-CoV-2 infection

A549-hACE2 lung carcinoma cells expressing the human ACE2 protein (Invivogen) were maintained in DMEM with 4.5 g/L glucose and 2 mM L-glutamine (Gibco), 10% heat-inactivated fetal bovine serum (FBS, R&D Systems), 100 U/mL penicillin, 100 µg/mL streptomycin (Gibco), and 0.5 µg/mL puromycin (Sigma). Vero E6 TMPRSS2 cells were maintained in DMEM with 4.5 g/L glucose and 2 mM L-glutamine (Gibco), 10% heat-inactivated FBS (R&D Systems), 100 U/mL penicillin and 100 µg/mL streptomycin (Gibco), and 5 µg/mL blasticidin (Invivogen). SARS-CoV-2 strain 2019n-CoV/USA-WA-1/2020 (BEI, Cat. #NR-52281) was propagated on Vero TMPRSS2 cells as described (Bohan et al., 2021). Infected cell supernatant containing virus was collected, passed through a 0.45 µM filter, and concentrated by centrifugation at 10,000 × g for 24 hours at 10°C. Virus titer was determined by TCID_50_ assay on Vero TMPRSS2 cells (Bohan et al., 2021).

### Immunofluorescence

At 24 hours post-infection, A549-hACE2 cells were fixed with 4% PFA (Electron Microscopy Sciences, Cat. #15710) for 30 min at room temperature and permeabilized with 0.5% Triton X-100 in phosphate buffered saline. Cells were incubated with rabbit monoclonal SARS-CoV/SARS-CoV-2 nucleocapsid antibody (SinoBiological 40143-R001, dilution 1:100), followed by incubation with Alexa Fluor 488 goat anti-rabbit IgG (Life Technologies, dilution 1:500). Nuclei are stained with DAPI (1 µg/ml). Images were acquired with a Nikon Eclipse Ts2 microscope.

### RT-qPCR

Total cellular RNA was purified using TRIzol reagent according to the manufacturer’s protocol (Invitrogen). RNA concentration was determined using a Nanodrop-OneC spectrophotometer (Thermo Fisher). RNA was DNase-treated using TURBO DNase according to the manufacturer’s protocol (Thermo). 100 ng DNase-treated RNA was used as template for cDNA synthesis using the iScript cDNA Synthesis Kit according to the manufacturer’s protocol (Bio-Rad). The resulting cDNA was diluted 1:10 and 2 µL was used for qPCR amplification of SARS-CoV-2 spike using iTaq Universal SYBR Green Supermix according to the manufacturer’s protocol (Bio-Rad). Copy number was calculated in reference to a standard curve of known copy number (10^2^ – 10^7^ copies spike). The following two primers were used at a final concentration of 200 nM (Bohan et al., 2021):

SARS-CoV-2 S forward: 5’ - CTACATGCACCAGCAACTGT – 3’

SARS-CoV-2 S reverse: 5’ - CACCTGTGCCTGTTAAACCA – 3’

### m^6^A immunoprecipitation

Total cellular RNA was purified using TRIzol reagent according to the manufacturer’s protocol (Invitrogen). RNA concentration was determined using a Nanodrop-OneC spectrophotometer (Thermo Fisher). 5 μg total RNA and m^6^A spike-in control mixture were added to 300 μL 1× IP buffer (50 mM Tris-HCl, pH 7.4, 150 mM NaCl, 0.1% NP40, 40U/μL RNase Inhibitor) containing 2 μg anti-m^6^A rabbit polyclonal antibody (Synaptic Systems). The reaction was incubated with head-over-tail rotation at 4°C for 2 hours. 20 µL Dynabeads™ M-280 Sheep Anti-Rabbit IgG suspension per sample was blocked with freshly prepared 0.5% BSA at 4°C for 2 hours, washed three times with 300 μL 1× IP buffer, and resuspended in the total RNA-antibody mixture prepared above. The RNA binding to the m^6^A-antibody beads was carried out with head-over-tail rotation at 4°C for 2 hours. The beads were then washed three times with 500 μL 1× IP buffer and twice with 500 μL Wash buffer (50 mM Tris-HCl, pH 7.4, 50 mM NaCl, 0.1% NP40, 40 U/μL RNase Inhibitor). The enriched RNA was eluted with 200 μL Elution buffer (10 mM Tris-HCl, pH 7.4, 1 mM EDTA, 0.05% SDS, 40U Proteinase K, 1 μL RNase inhibitor) at 50°C for 1 hour. The RNA was extracted by acid phenol-chloroform and ethanol precipitated.

### RNA labeling and hybridization

The immunoprecipitated (IP) RNAs and unbound RNAs were mixed with equal amount of calibration spike-in control RNA, separately amplified and labeled with Cy3 (unbound) and Cy5 (IP) using Arraystar Super RNA Labeling Kit. The synthesized cRNAs were purified by RNeasy Mini Kit (Qiagen). The concentration and specific activity (pmol dye/μg cRNA) were measured with NanoDrop ND-1000. Cy3 and Cy5 labeled cRNAs were mixed and 2.5 µg of the cRNA mixture in 19 µL volume was fragmented by adding 5 μL 10× Blocking Agent and 1 μL of 25× Fragmentation Buffer, followed by heating at 60°C for 30 min. The fragmented RNA was combined with 25 μL 2× Hybridization buffer. 50 μL hybridization solution was dispensed into the gasket slide and assembled to the m^6^A-mRNA&lncRNA Epitranscriptomic Microarray slide. The slides were incubated at 65°C for 17 hours in an Agilent Hybridization Oven. The hybridized arrays were washed, fixed, and scanned using an Agilent Scanner (G2505C).

### Data and statistical analysis

Agilent Feature Extraction software (version 11.0.1.1) was used to analyze acquired array images. Raw intensities of IP (Cy5-labelled) and unbound (Cy3-labelled) were normalized using the average of log2-scaled Spike-in RNA intensities. After Spike-in normalization, the probe signals having present or marginal QC flags in at least 3 out of 6 samples were retained for further “m^6^A quantity” analyses. “m^6^A quantity” was calculated for the m^6^A methylation amount based on the IP (Cy5-labelled) normalized intensities. Differentially m^6^A-methylated RNAs between two comparison groups were identified by filtering with the fold change and statistical significance (p-value) thresholds. Protein-coding transcripts that were found to be differentially m^6^A-methylated between mock and SARS-CoV-2-infected samples with a fold change ≥1.5 and a p-value ≤0.05 were uploaded to iPathwayGuide (ipathwayguide.advaitabio.com) for gene ontology, pathway, upstream regulator, and network analyses (Ahsan and Draghici, 2017; Donato et al., 2013; Draghici et al., 2007; Tarca et al., 2009).

## CRediT authorship contribution statement

**Stacia Phillips:** Conceptualization, Methodology, Formal analysis, Investigation, Writing – Original Draft, Writing – Review & Editing, Visualization, Supervision, Project administration. **Shaubhagya Khadka:** Investigation. **Dana Bohan:** Investigation, Writing – Review & Editing. **Constanza Espada:** Investigation, Writing – Review & Editing. **Wendy Maury:** Methodology, Resources, Writing – Review & Editing, Supervision. **Li Wu:** Conceptualization, Methodology, Writing – Review & Editing, Visualization, Supervision, Project administration, Funding acquisition.

## Acknowledgments

The following reagent was deposited by the Centers for Disease Control and Prevention and obtained through BEI Resources, NIAID, NIH: SARS-Related Coronavirus 2, Isolate USA-WA1/2020, NR-52281. We thank Dr. Michael Chimenti for helpful discussion and assistance with iPathway Guide analysis. We thank Dr. Patrick Sinn for A549-hACE2 cells.

## Supplementary Tables (two separate Excel files)

**Table S1:** All differentially m^6^a-modified transcripts in SARS-CoV-2-infected A549-hACE2 cells vs. mock-infected control cells.

**Table S2:** All pathways associated with at least one differentially methylated mRNA in SARS-CoV-2-infected A549-hACE2 cell vs. mock-infected control cells.

